# High-resolution global peptide-protein docking using fragments-based PIPER-FlexPepDock

**DOI:** 10.1101/174714

**Authors:** Nawsad Alam, Oriel Goldstein, Bing Xia, Kathryn A. Porter, Dima Kozakov, Ora Schueler-Furman

**Affiliations:** Department of Microbiology and Molecular Genetics, Institute for Medical Research Israel-Canada, Hadassah Medical School, The Hebrew University, Jerusalem 91120, Israel; School of Computer Sciences and Engineering, The Hebrew University, Jerusalem 9190416, Israel; Department of Biomedical Engineering, Boston University, 44 Cummington Street Boston, MA 02215, USA; Department of Applied Mathematics and Statistics, Stony Brook University, Stony Brook, NY 11794; Laufer Center for Physical and Quantitative Biology, Stony Brook University, Stony Brook, NY 11794; Institute for Advanced Computational Sciences, Stony Brook University, Stony Brook, NY 11794

## Abstract

Peptide-protein interactions contribute a significant fraction of the protein-protein interactome. Accurate modeling of these interactions is challenging due to the vast conformational space associated with interactions of highly flexible peptides with large receptor surfaces. To address this challenge we developed a fragment based high-resolution peptide-protein docking protocol. By streamlining the Rosetta fragment picker for accurate peptide fragment ensemble generation, the PIPER docking algorithm for exhaustive fragment-receptor rigid-body docking and Rosetta FlexPepDock for flexible full-atom refinement of PIPER docked models, we successfully addressed the challenge of accurate and efficient global peptide-protein docking at high-resolution with remarkable accuracy. Validation on a representative set of solved peptide-protein complex structures demonstrates the accuracy and robustness of our approach, and opens up the way to high-resolution modeling of many more peptide-protein interactions and to the detailed study of peptide-protein association in general. PIPER-FlexPepDock is freely available to the academic community as a server at http://piperfpd.furmanlab.cs.huji.ac.il.

## Introduction

Proteins are the workhorses inside living cells, and interactions among them are critical for various important biological processes ^1^. A significant fraction of these interactions (15-40%) ^2^ are peptide mediated, where a short stretch of residues from one partner contributes most to its binding to the other. Such short peptidic regions, also termed short linear interacting motifs (SLIMs) are often found embedded inside disordered regions of intrinsically disordered proteins (IDPs) ^2^,^3^, or appear as flexible linkers connecting domains ^4^ and as flexible loops tethered to rigid segments ^5^.

The development of accurate structure based modeling tools is critical for atomic level understanding of peptide-protein interactions, to allow the manipulation of known interactions, to discover yet unknown peptide-protein interactions and networks, and to provide starting points for the design of novel peptides and related molecules to target specific systems of pharmacological interest ^6^. A number of computational tools have been developed to assist the characterization of peptide-protein interactions, including the prediction of peptide binding sites ^7^-^9^, refinement of coarse peptide-protein models 10, folding and docking on a known binding site ^11^ and most challenging of all, global peptide-protein docking with no prior information about the peptide structure and the binding site ^12^-^17^. The challenges associated with the global docking of flexible peptides have been addressed in different ways, by reducing the conformational space to be sampled both for the internal degrees of freedom of the peptide as well as its rigid-body orientations on the receptor surface. For peptide docking within the HADDOCK docking framework ^12^, the peptide backbone is represented by idealized conformation(s), such as alpha helix, beta strand and polyproline-II, followed by rigid-body, semi-flexible and fully-flexible docking with explicit solvation ^18^. The pepATTRACT protocol ^13^,^19^ uses the same approach to represent the peptide, followed by coarse-grained rigid-body docking and flexible full-atom refinement. The AnchorDock protocol uses molecular dynamics simulations to generate a set of plausible peptide conformations, which are then docked using anchor-driven simulated annealing molecular dynamics around predicted anchoring spots on the receptor ^14^. The CABS-dock protocol uses randomly generated peptide conformations based on either predicted or known secondary structure, randomly orients these peptides over the receptor surface, and refines them using replica exchange Monte Carlo dynamics ^15^. The MDockPep protocol ^16^ uses peptide sequence similarity to extract fragments from high resolution protein structures, which are further refined using MODELLER ^20^ to generate plausible peptide conformations, and then docked onto the receptor using rigid-body docking and flexible docking with AutoDock Vina ^21^. The recently published IDP-LZerD protocol models the binding of long disordered segments to structured proteins using the Rosetta fragment picker protocol ^22^ to generate fragments of 9-residue overlapping windows followed by LZerD 23 rigid-body docking and molecular dynamics refinement ^17^. Finally, we have recently advanced a novel, global motif-based peptide fragment docking approach, PeptiDock ^24^, in which peptide binding motif information rather than secondary structure propensity is used to extract fragments from the Protein Data Bank (PDB ^25^), which are then docked to the receptor using PIPER rigid body docking ^26^, followed by minimization using CHARMM ^27^.

These significant recent advances in global peptide docking notwithstanding, present approaches are still limited in their modeling quality and general applicability, and there is ample room for improvements that would enable the detailed high-resolution study of more peptide-protein interactions with higher accuracy. Here we describe PIPER-FlexPepDock, a successful effort toward the development of such a robust, highly accurate, global peptide-protein docking protocol. By integrating accurate peptide fragment ensemble generation using the Rosetta fragment picker ^22^, fast and exhaustive fragment-receptor rigid-body docking using PIPER docking ^28^, and flexible full-atom refinement of coarse PIPER models using Rosetta FlexPepDock ^10^, we were able to sample both the peptide backbone conformational states, as well as the landscape of the peptide-receptor interactions very efficiently and with much higher accuracy than current protocols: on a non-redundant dataset of peptide-protein complexes (**Table 1** below), PIPER-FlexPepDock generates for about half models within 2.5 Å ligand RMSD (2.0 Å, when restricted to motif regions where available), more than twice as many as for existing peptide docking protocols such as pepATTRACT ^13^ (among the 10 top-ranked predictions; **Table 2** below).

Our results highlight the relevance of representing the peptide as a set of fragments that can be exhaustively docked as rigid bodies onto the receptor structure and subsequently refined using an accurate refinement protocol. They reinforce the underlying biophysical model of a conformer ensemble of the free peptide that already samples the bound conformation (at least in the encounter-complex, protein-like environment) and involves only limited induced fit, not unlike to the classical association between preformed protein domains. As a result, PIPER-FlexPepDock brings into reach the study and targeted manipulation of a range of additional peptide-mediated interactions not accessible before due to limitations in sampling and/or accuracy.

## Results

### Overview of the PIPER-FlexPepDock protocol (Figure 1)

**Step A | Generation of fragment set to represent the peptide conformer ensemble:** In a previous study we have shown that the bound peptide conformation can be well represented by extraction of short fragments from the PDB based on information of known binding sequence motifs ^24^. Here we have generalized this approach beyond motifs, using fragment libraries selected by the Rosetta fragment picker protocol ^22^ based on sequence and secondary structure similarity (see **Methods**). The coordinates of the top 50 mapped fragments are extracted from the PDB, including both backbone and side-chain atoms, and non-identical residues in the extracted fragments are mutated to the desired sequence. This set of fragments adequately represents the peptide conformational ensemble, sampling also its receptor bound conformation (see below). The peptide may be trimmed in cases where information is available about the range of the binding segment (from motif databases such as the Eukaryotic Linear Motif (ELM) resource ^29^,^30^, literature, or experiments such as alanine scanning), since fragments generated for shorter peptide sequences are usually better representative than longer fragments (as, e.g., for loop modeling ^31^), and fraying ends beyond the motif may contribute less to determine critical binding details.

### Step B | Fragment rigid-body docking using PIPER

Each of the fragments is docked onto the receptor structure using PIPER, an exhaustive Fast Fourier Transform (FFT)-based rigid body docking algorithm ^28^, as implemented previously for PeptiDock ^24^ (see **Methods**), and top ranking fragment orientations from each docking run are collected and combined together. These models are of low resolution as no flexibility is included in the PIPER algorithm, and therefore ranked using a soft potential that allows a certain degree of steric clashes to overcome the limitations of rigid-body only docking.

**Figure 1.**
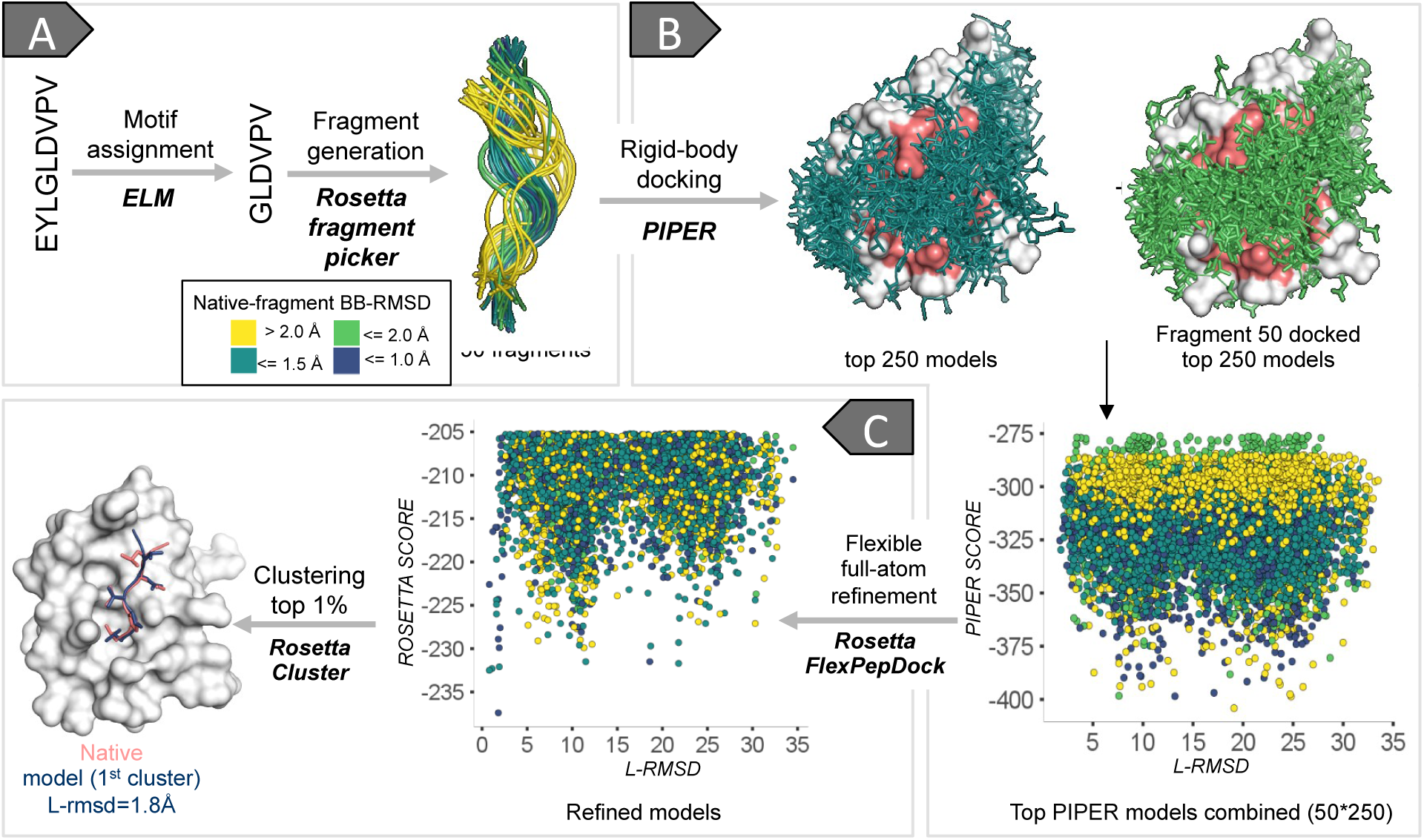
Overview of the PIPER-FlexPepDock peptide docking protocol. Example shown: PDZ domain-peptide interaction [PDB IDs of receptor structure 1MFG (bound) and 2H3L (free)]. For a given receptor structure and peptide sequence, the divide and conquer strategy involves first the description of the peptide as an **ensemble of fragments (A)**, their fast and exhaustive **rigid body docking** (using PIPER) onto the whole receptor (binding site region is shaded salmon) **(B)**, and subsequent **high-resolution refinement** (using Rosetta FlexPepDock; the top 5000 models are included in the plot) **(C)**, followed by clustering and selection of top ranking representatives. Fragments are colored according to their similarity to the native bound peptide conformation. L-RMSD: Ligand root mean square deviation from crystal structure; see text for more details.

### Step C | FlexPepDock refinement of PIPER models and selection of final models

Each of the PIPER models is refined by a single fully flexible refinement run using the Rosetta FlexPepDock Refinement algorithm ^10^ (see **Methods**). The top ranking refined models are clustered (as in Gray *et al*. ^32^), clusters are ranked based on the reweighted score of the best scoring model in each cluster (as in Raveh *et al*. ^11^), and the top 10 ranked cluster representatives are selected as prediction (following the CAPRI scheme that accepts 10 models ^33^).

### Initial calibration of the PIPER-FlexPepDock on a small set of protein-peptide complexes

Motivated by our recent advance in global peptide docking using a motif-focused approach ^24^ we ventured into the development of a more generalized protocol. We initially calibrated our docking approach on a small representative set of nine peptide– protein complexes (highlighted in bold in **Table 1**; see also **Supplementary Table S1A**). We trimmed the peptide based on the motif defined in ELM, where available. For all complexes impressive modeling accuracy was achieved for this new global docking approach (within ≤2.5Å Ligand RMSD models among the top 10 ranking clusters; **Table 1**). For the full length peptides modeling near-native models were obtained for 5/9 cases, highlighting the benefits for motif (or shorter peptide sequence) focused modeling, due to better fragment quality compared to the corresponding full-length peptides (**Table 1**). Encouraged by these initial results, we proceeded to the validation of our protocol on a larger and representative set of peptide-protein complexes (**Table 1** and **Supplementary Table S1B**).

**Table 1.**
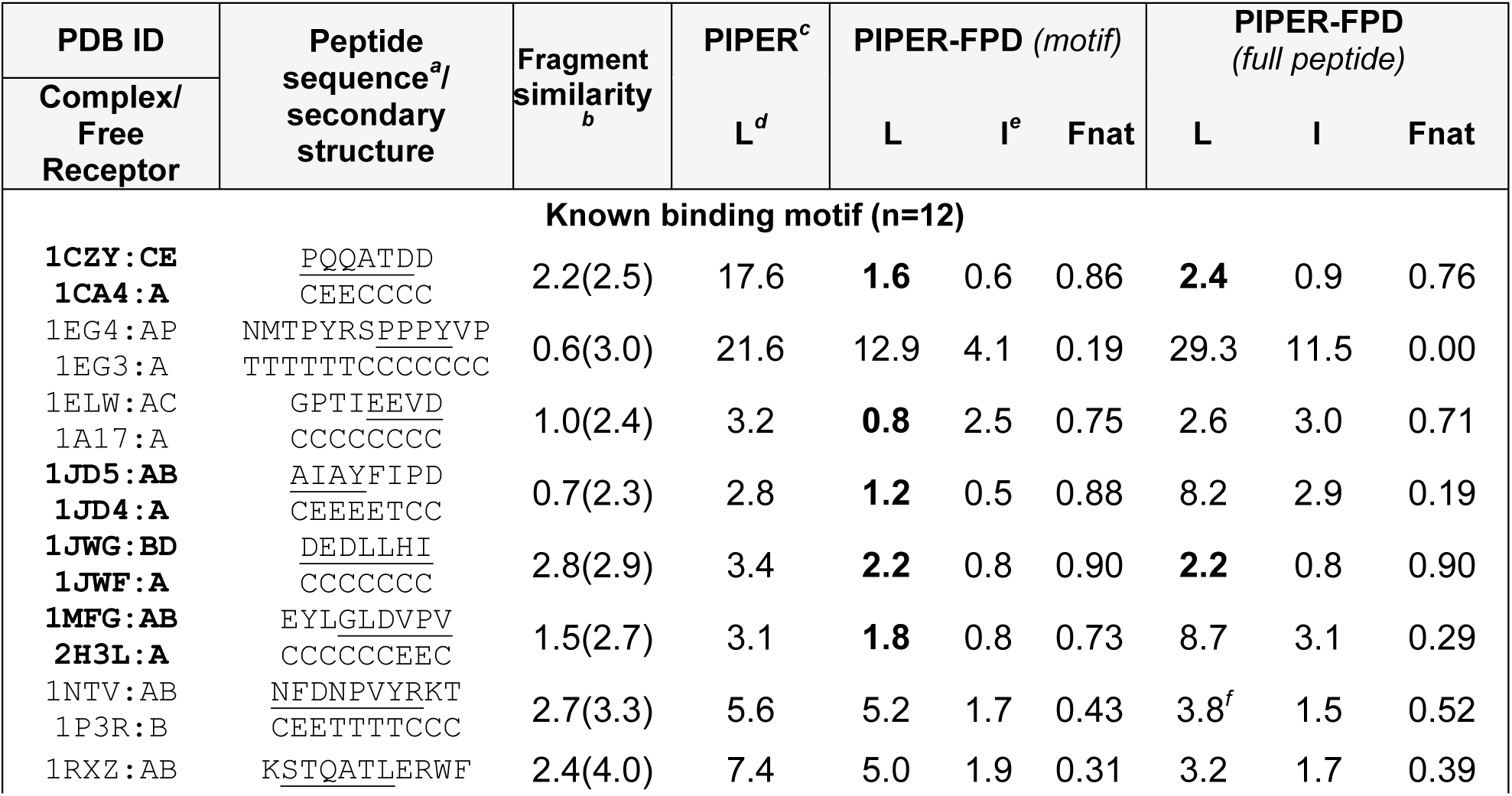
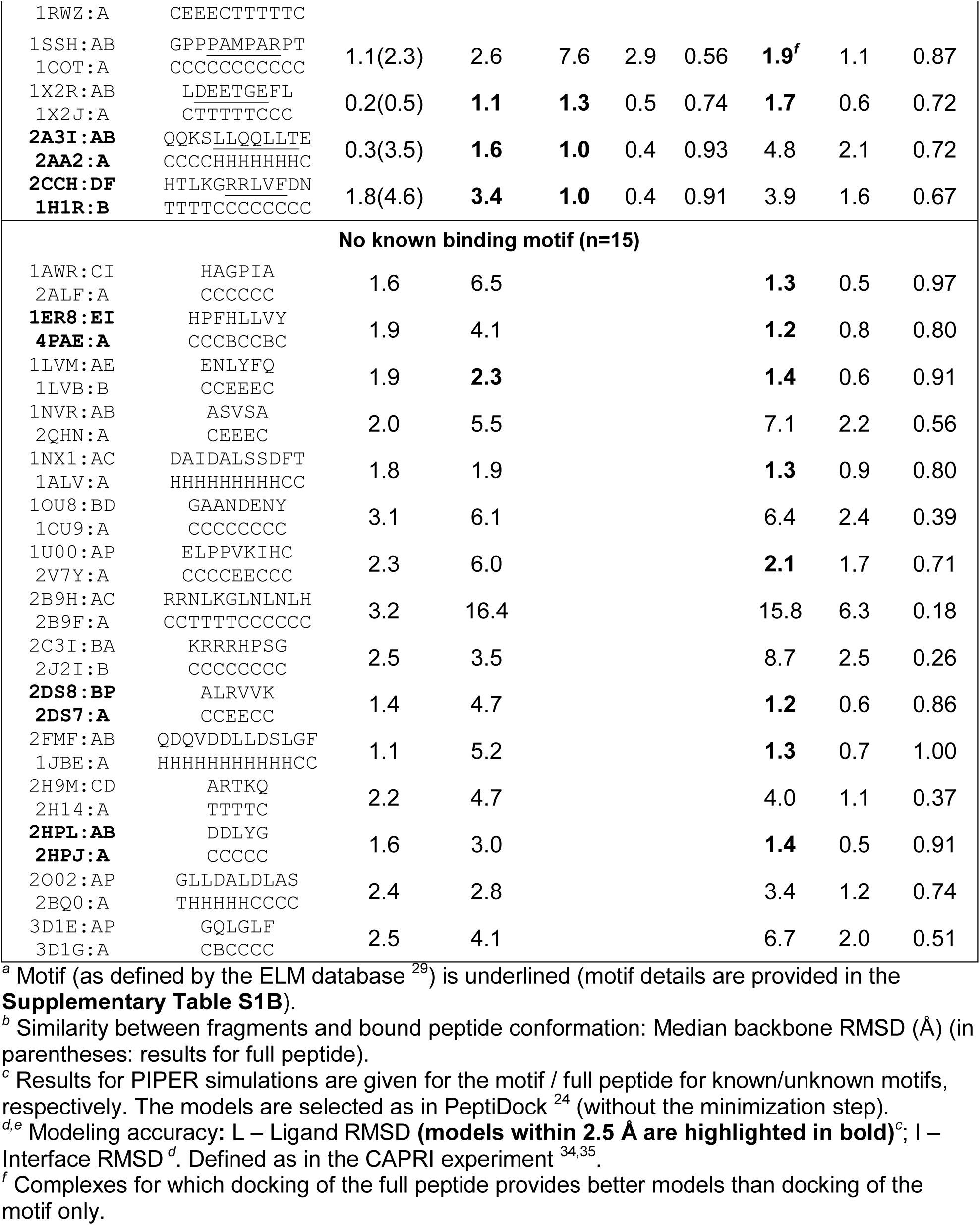
Benchmark of peptide-protein complexes used in this study (non-redundant set; see **Table S1C** for full set). PDB ids of the initial calibration set are highlighted in bold.

### Assessment of peptide docking performance

We assessed the performance of PIPER-FlexPepDock on a larger, non-redundant set consisting of 27 complexes (non-redundant at the domain level, as defined by CATH ^36^; see **Methods**), among them 12 with reported binding motif. The benchmark is summarized in **Table 1** (**Supplementary Table S1C** provides results for a redundant set of 42 complexes used in previous studies, as well as additional details, including performance of other approaches for comparison).

### Representing the peptide conformational states using fragments

Fragments derived from solved protein structures contain valuable information about the local structural context that can be used to efficiently reduce sampling space for various modeling applications, including e.g. *ab initio* protein folding ^37^ and loop building ^31^,^38^,^39^. In our protocol we use the Rosetta fragment picker protocol ^22^ to generate fragments consistent with both the peptide sequence and the (predicted) secondary structure (See **Methods**).

How accurately do the fragments represent the peptide conformational states? Most importantly, how similar are peptide conformations to the one adapted when bound to their receptor? A significant representation of similar fragments could guarantee that, when docked with high density in the binding site using an exhaustive but accurate rigid-body docking algorithm, they could efficiently be refined to high resolution using an accurate refinement algorithm such as Rosetta FlexPepDock. To assess the quality of the fragments (*i.e.,* their coverage of the bound conformation) we analyzed the distribution of backbone RMSDs of the fragments relative to the bound peptide conformation. Reassuringly, the fragment pool generated using the Rosetta fragment picker protocol represents the bound like peptide conformation with high accuracy in our benchmark of 27 peptide-protein complexes (**Figure 2A**: median backbone RMSD within 2.0 Å for 15 out of the 27 cases, with average backbone RMSD of the best ten fragments within 1.0 and 1.5 Å for 14 and 21 cases, respectively). The best accuracy is achieved for helical peptide motifs (e.g., the helical nuclear receptor box motif in 2A3I ^40^; for helical peptides with coiled terminus segments such as 2FMF ^41^ and 1NX1 ^42^ the median backbone RMSD is slightly higher). Even for the remaining peptides the fragment ensemble is often composed of a significant portion of bound like representatives. The worst representation of bound-like peptides is obtained for few longer coil peptides, such as 2B9H ^43^, which defines the limitation of the fragment picker protocol for longer sequences. In such cases, trimming the peptide might improve the quality significantly.

**Figure 2.**
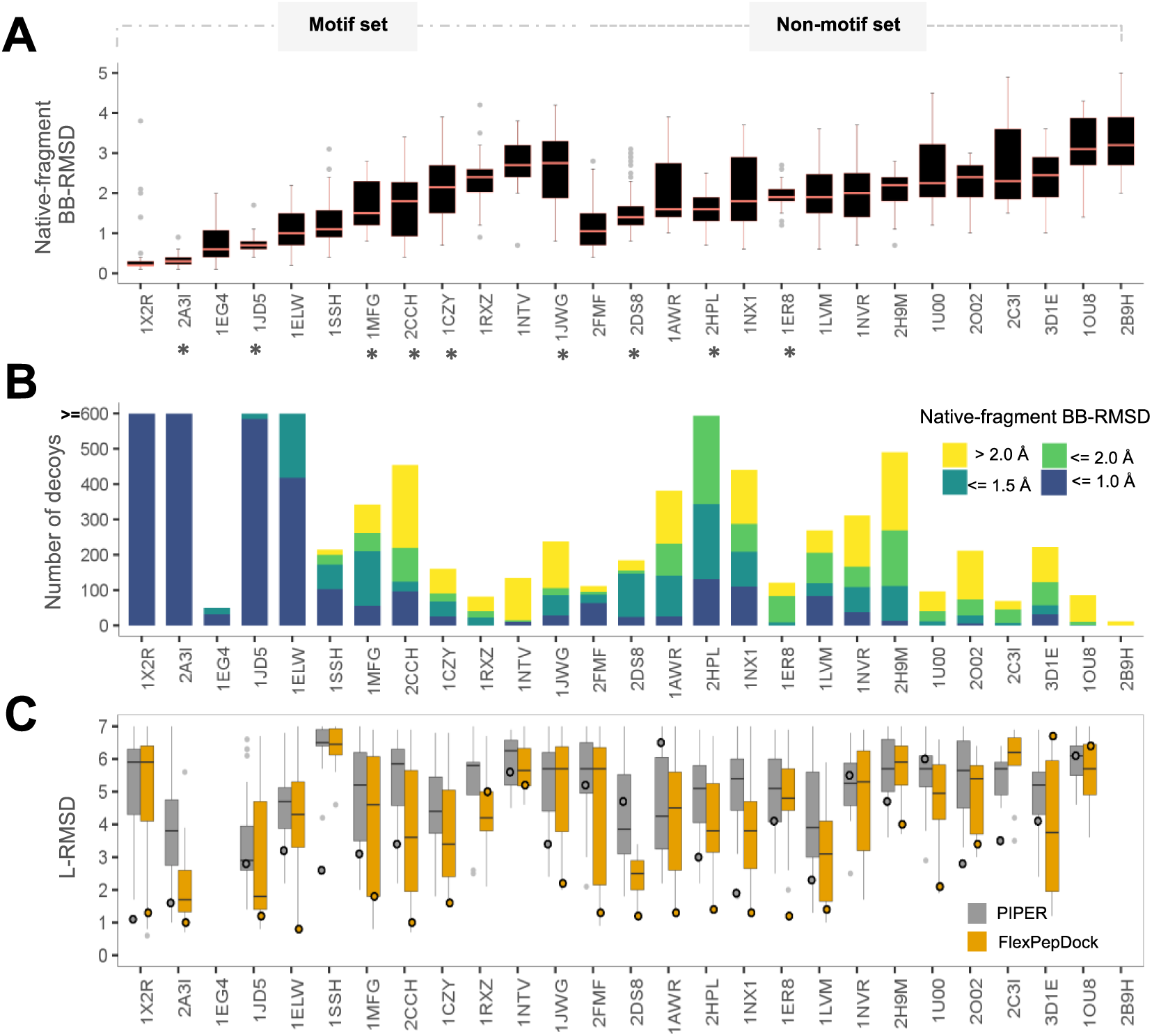
Assessment of performance of the different steps of PIPER-FlexPepDock. (A) Fragment quality: distribution of fragment backbone RMSDs relative to the native bound peptide conformation (defined as fragment quality). PDBs with and without motif information are grouped separately. The initial calibration set is marked with asterisks (*). **(B) PIPER rigid body docking:** distribution of the number of models within 5Å ligand (L)-RMSD from the native, colored according to fragment quality**. (C) Improvement after FlexPepDock refinement:** distribution of the L-RMSDs of the top 1% FlexPepDock refined models (in orange) and corresponding PIPER models (in gray). Shown are the results of runs starting from the unbound receptor structure and including receptor minimizations (see also **Figure 3**). The circles represent the L-RMSD values of the best model among the top 10 ranking clusters. The Y-axis has been trimmed to 7Å. (Note that the gray circles are taken from the top-ranked models of the *full* PIPER run based on density clustering [see **Methods** and Porter *et al*. ^24^]), while the distributions represent the subset of models that served as starting structures for the models selected after FlexPepDock refinement).

We previously showed that extracting fragments based on sequence motif information allows identification of bound peptide conformations that reflect the structural pattern of these motifs ^24^. We demonstrate here that representative fragments are not restricted to peptides with known motifs. In fact, a comparison to the fragments extracted based on sequence motif (for the dataset analyzed in the PeptiDock study, using the motif definition therein ^24^) shows that the fragment picker approach produces overall ensembles that contain structures more similar to the bound peptide conformation (see **Supplementary Table S2**).

### Rigid-body docking: fragment quality and PIPER performance

The fact that the fragment ensembles include bound-like conformations justifies proceeding to the next step, namely their docking onto the receptor. The PIPER rigid-body docking protocol allows fast and exhaustive sampling to provide coarse models of fragment-receptor interactions, of which the top-scoring can be followed up by subsequent refinement to allow for conformational changes upon binding. The effective range for successful refinement using the FlexPepDock protocol was previously found to be within up to 5Å in terms of Cartesian RMSD, and up to 50 degrees in terms of ϕψ RMSD (distance of fragment from the bound peptide conformation in ϕψ dihedral space) ^10^. It is thus important for the PIPER docking stage to identify a large pool of fragments that are densely docked in close proximity (within effective Cartesian RMSD range) of the native peptide binding mode, involving docked fragments that are similar to the native peptide bound conformation (within effective phi-psi RMSD range). Indeed, analysis of the top ranking PIPER models shows presence of good quality fragments at the binding site (in fact, most complexes include <1.0Å bb RMSD fragments; **Figure 2B**).

### Improvement of PIPER models by FlexPepDock refinement

The FFT algorithmic implementation of rigid-body sampling in PIPER makes exhaustive orientation search possible with significant computational efficiency, but is defined on a grid. Consequently, the scoring function can successfully isolate the best few hundreds from the vast pool of billions of positions of the peptide fragment relative to the receptor, but not discriminate the top rigid-body docked models further (**Figure 3A** & **Supplementary Figure S1**). In turn, the Rosetta scoring function used in the FlexPepDock Refinement protocol (currently Talaris 2014 ^44^) is highly accurate, but this flexible docking protocol lacks the ability for fast and exhaustive sampling. Thus, to address the problem of exhaustive sampling with high accuracy, we combine the fast and exhaustive rigid-body sampling of PIPER with accurate flexible refinement by FlexPepDock of the top ranking few hundred best models. Indeed, the FlexPepDock refinement stage significantly improves the model quality, as well as better model ranking (See **Figures 2C, 3C** & **Supplementary Figure S1**). This includes the identification of a near-native funnel missed before (e.g. 1CZY in **Figure 3 – compare A** to **C**), or significant enhancement of a near-native funnel (e.g.1JD5 and 2A3I). More examples can be found in **Supplementary Figure S1**.

We performed three runs to assess protocol performance (Summarized in **Figure 4A** and **Supplementary Table S1B;** specific examples are shown in **Figure 3**): First, we applied the protocol to bound receptor structures. For these runs a near-native peptide conformation (L-RMSD <= 2.0Å, see **Methods** section) was found among the top 10 ranked clusters for 19 out of 27 complexes (success rate=70%**, Figure 3D**). We then proceeded to the real-world scenario, in which the free receptor structure was provided as starting point (unbound run), leading to worse performance, as expected (10 complexes successfully modeled - success rate=37%, **Figure 3B**). Importantly however, when including also receptor flexibility during the refinement stage (unbound-min run), these results improved, in particular if 10 best models are considered (14/27 complexes successfully modeled - success rate=52%, **Figure 3C**).

**Figure 3.**
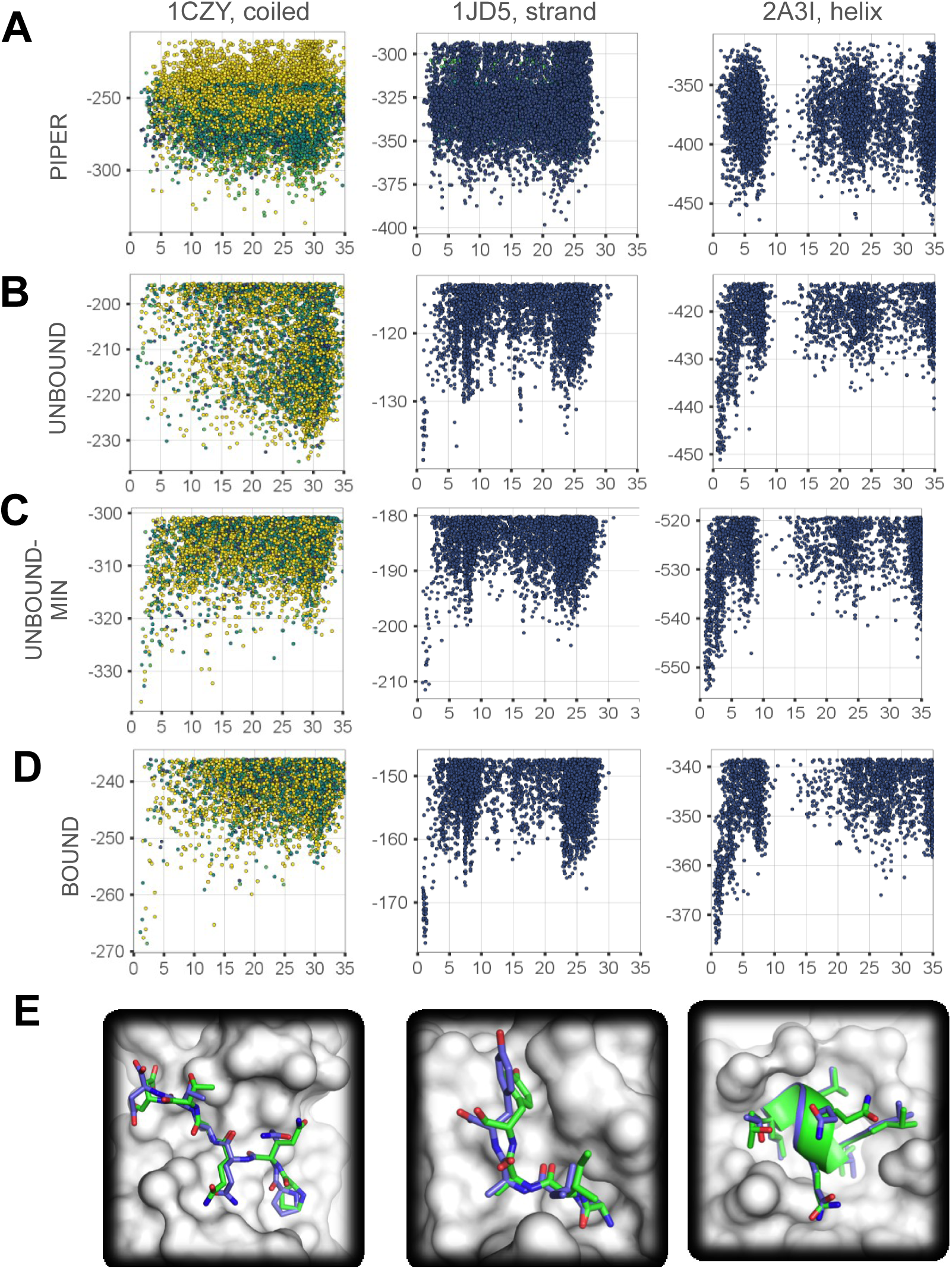
Examples of global peptide docking energy landscapes: Left: PDB id 1CZY (coiled peptide); Center: 1JD5 (extended peptide); Right: 2A3I (helical peptide). (A) Energy landscape as sampled in the first docking step of the protocol by PIPER rigid body docking of peptide fragments onto the *unbound* receptor structure. **(B-D)** Energy landscapes for the PIPER-FPD scheme, starting from the unbound receptor structure **(B**), the unbound receptor structure including receptor flexibility **(C)**, and the corresponding bound receptor for comparison **(D).** Models are colored according to fragment quality, as in previous Figures. **(E)** Comparison of the modeled to the native structure (shown in blue and green, respectively).

**Figure 4.**
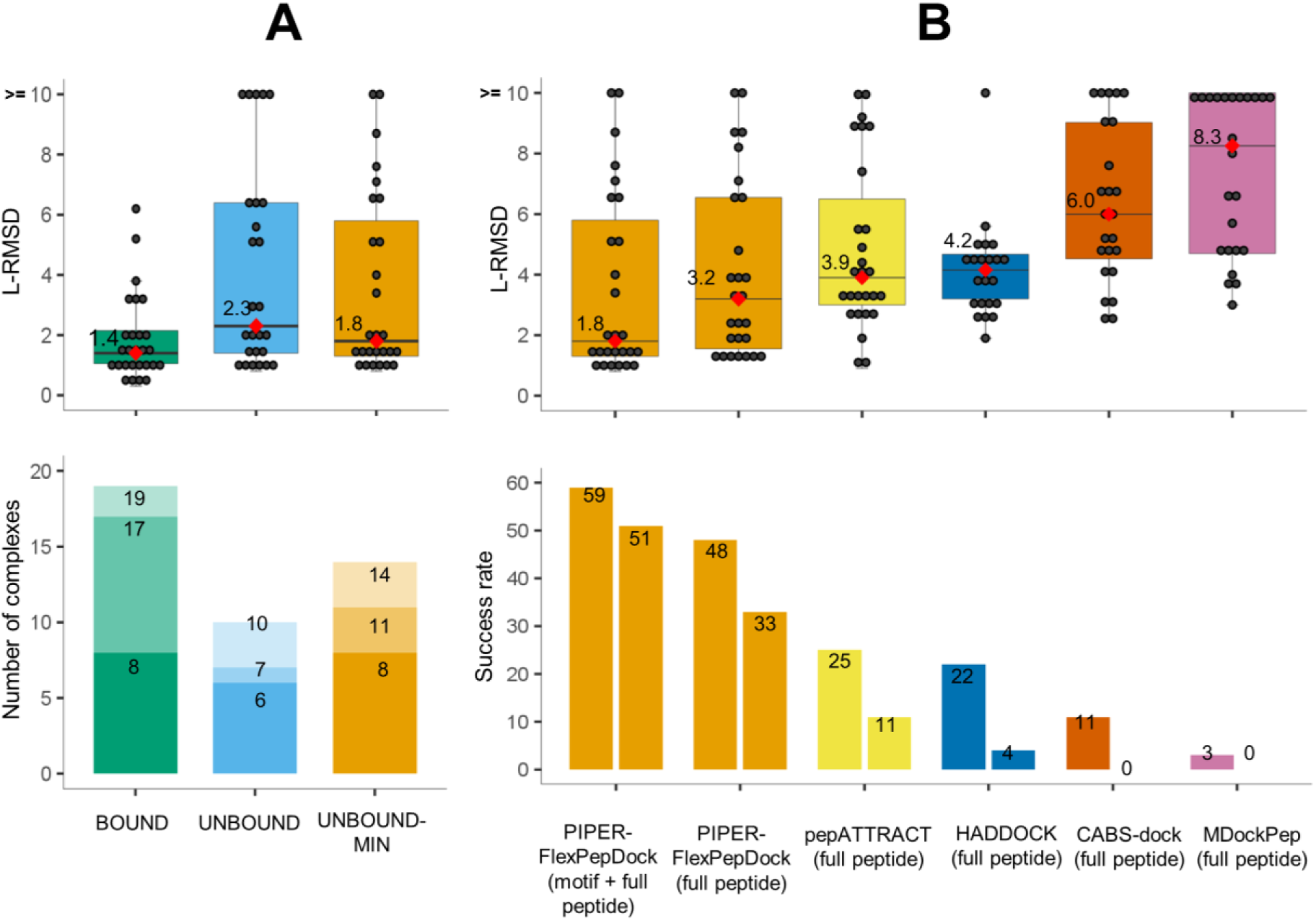
PIPER-FlexPepDock peptide docking performance. **(A) Overall performance on a non-redundant set of 27 peptide-protein complexes. Top:** Distribution of best model L-RMSDs (among top 10 ranking clusters) for runs using the bound (BOUND) and free (UNBOUND & UNBOUND-MIN) receptor structures, the latter including also receptor flexibility in the final refinement step (only the motif region was modeled for the 12 complexes with known motif). The median values are shown as red diamonds and printed alongside. **Bottom:** Distribution of the ranks of the first near-native cluster (defined as L-RMSD <=2.0Å), shown using different shades (for corresponding results among the top1, top3 and top10 ranked predictions)**. (B**) Comparison to performance by other algorithms**. Top:** Box plots of best L-RMSDs among top 10 ranking clusters, including results for the motif part where the motif is known (as in A), as well as for the full peptide, for comparison. **Bottom:** Performance **i**s shown for different cutoffs (2.0Å and 3.0Å L-RMSD in left and right boxes, respectively) (See **Supplementary Table S1B** for more details).

### Comparison with other global docking protocols

We compared the results of PIPER-FlexPepDock (unbound-min run) with other existing global peptide-protein docking protocols such as HADDOCK ^12^, pepATTRACT ^13^, CABS-dock ^15^, and MDockPep ^16^ on our non-redundant set of 27 complexes, as well as on the set of 42 complexes used by these protocols in previous studies (34 complexes were compared with HADDOCK as other 8 cases were not included in their unbound run set). Since full length peptides were modeled using the other protocols, we modeled full length peptides for the motif set cases for valid comparison. The success rate for generating near-native models (*i.e.,* L-RMSD within 2.0Å, or 3.0Å) was significantly better for PIPER-FlexPepDock than any other protocol, even for models of the full peptides (see **Figure 4B** and **Table 2**).

**Table 2.**
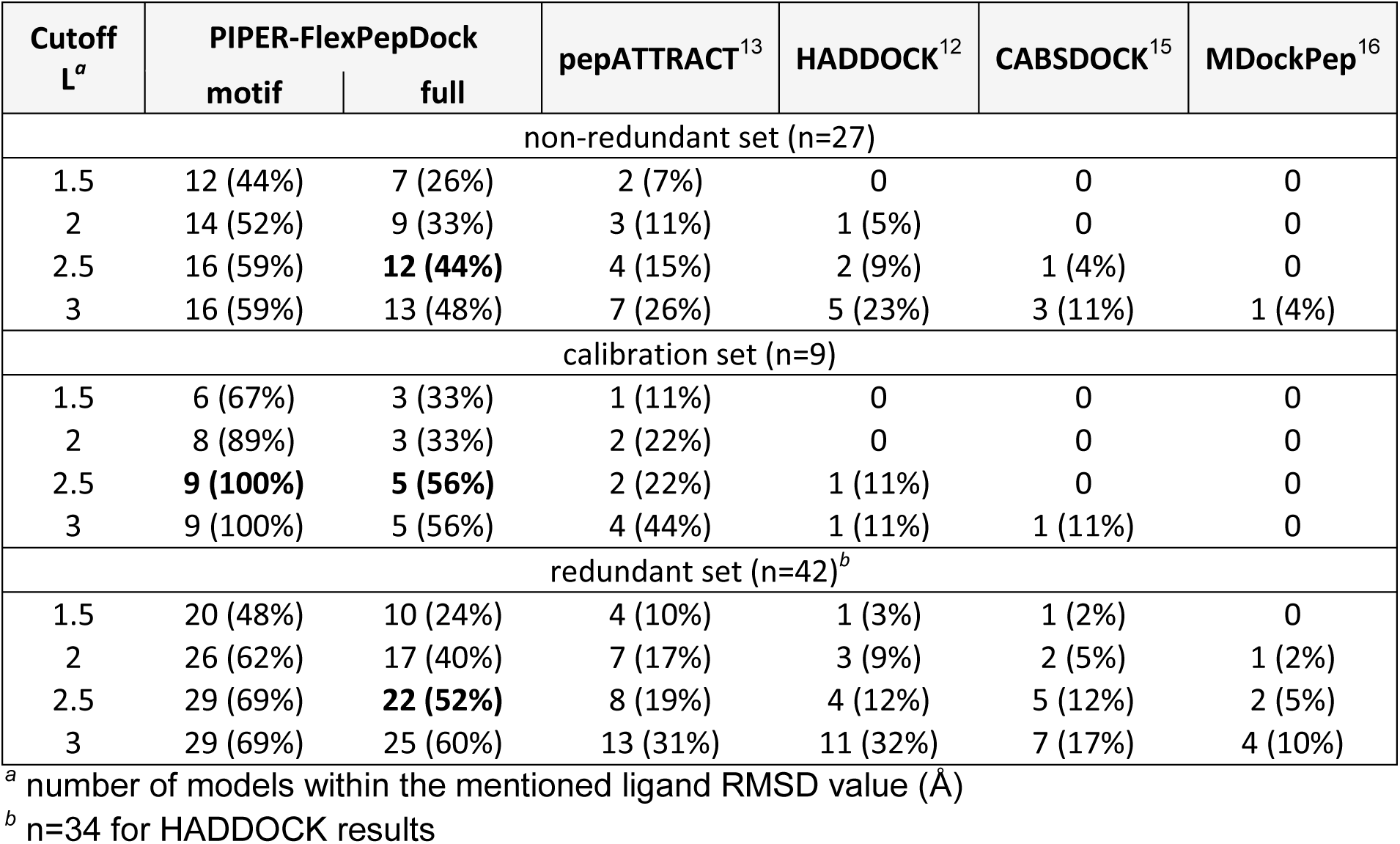
Summary of performance of PIPER-FlexPepDock, and comparison to other peptide docking protocols. Results are shown for PIPER-FlexPepDock runs on unbound receptor structures, including receptor minimization.

### The PIPER-FlexPepDock server for the high-resolution modeling of peptide-protein interactions

In order to maximize the impact of our new protocol for global peptide-protein docking and to make it accessible to the modeling of many new peptide-protein complexes, we have set up a user-friendly server open to the scientific community (**Figure 5).** All that is needed is a structure of the receptor and a sequence of the peptide, but additional information about peptide secondary structure can also be included to narrow the search. The top-ranking resulting models can be downloaded, or inspected by an interactive viewer using the 3Dmol.js libraries ^45^.

**Figure 5.**
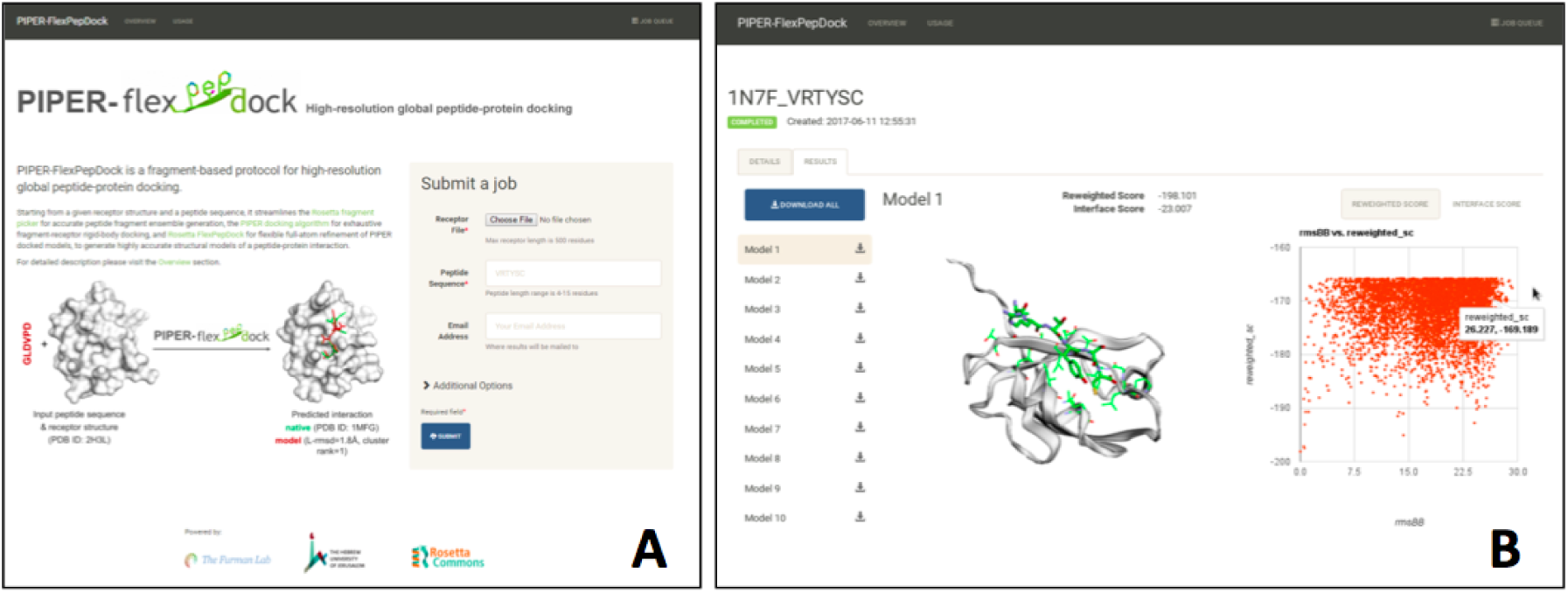
The PIPER-FlexPepDock server: (A) Job submission page: the required input includes the structure of the receptor and the sequence of the peptide; advanced options are accessible via a button. The tabs at the top provide links to detailed descriptions of the server, as well as to the Queue (upper right). **(B)** Results of an example peptide docking run: The liprin C-terminal peptide sequence VRTYSC docked onto the PDZ domain of GRIP1 (free receptor PDB id 1N7E). The top10 ranking models can be downloaded, and links to the individual models are provided to the left for inspection using an interactive viewer. In this case, Model 1 is an accurate prediction (L-RMSD=1.0Å from solved structure PDB id 1N7F). On the right side a scatter plot shows the sampled energy landscape (relative to an arbitrary reference structure).

## Discussion

### A new approach for global peptide docking with excellent performance *-*

With the presentation of our new PIPER-FlexPepDock algorithm, we have demonstrated that combining fast and exhaustive rigid-body docking (using the FFT-based PIPER docking algorithm) of a representative peptide conformer ensemble (approximated by fragments extracted from solved structures, based on local similarity of sequence and secondary structure), with high-resolution refinement (using Rosetta FlexPepDock) is a widely-applicable approach for the generation of models of peptide-receptor structures of remarkable accuracy – significantly better than any other current protocol - starting from the sequence of the peptide and the structure of the receptor. The performance on a large benchmark of solved peptide-protein complex structures demonstrates both accuracy and robustness of our modeling approach, and opens up the way of modeling many more peptide-protein interactions at much higher resolution and accuracy than any existing global peptide-protein global docking protocol.

### Receptor-bound peptide conformations are adequately represented by fragments extracted from protein monomer structures

This study demonstrates that fragments derived from solved protein structures, based on secondary structure and sequence similarity (rather than on sequence binding motifs which are not always available) represent the peptide conformational states with high accuracy, in particular the bound state. Interestingly, it is this same observation regarding the representation of local conformational preference that provided originally the platform for the breakthrough of Rosetta *ab initio* protein structure prediction ^46^. This indicates that while isolated peptides in solution rarely show significant conformational preferences ^47^, in the encounter complex regime in vicinity of other proteins, their conformational freedom seems to be restricted significantly (similar to local peptide regions within a full protein) and can be represented by fragment libraries, in concordance with previous reports that show similar arrangements of fragments within monomers and peptide-protein interactions ^48^.

### Effective sampling of the energy landscape

The simplified scoring function and exhaustive sampling with PIPER allows uniform sampling of the fragments onto the receptor on a smoothened energy landscape. The top scoring PIPER models represent the dense sampling into wider energy basins. Though the ranking of models might lack the accuracy at this stage, the following refinement stage performs local sampling to efficiently locate the minimum. Interestingly, this approach is much more effective than the local refinement starting from one representative model (only one FlexPepDock optimization run is necessary starting from each PIPER model, compared to several hundred to thousand runs starting from a representative (defined, e.g. from a PIPER run as implemented in the PeptiDock peptide motif docking algorithm ^24^). This is most probably due to the fact that these starting coarse models are trapped in many distinct states, each near a distinct local minimum, simplifying sampling during optimization.

### Mapping encounter complexes and more

The peptide-receptor binding energy landscape can provide a broader understanding of the binding mechanism itself. The exhaustive sampling with accurate refinement provides a high-resolution map of the energy landscape and helps us understand the energetic of the encounter between the peptide and the receptor. In a previous study, we were able to demonstrate that experimentally observed encounter complexes are well reproduced from a global protein docking energy landscape ^49^, and we anticipate that the corresponding peptide-protein docking energy landscape will provide similar information.

Our novel global peptide docking pipeline allows high-resolution modeling of peptide-protein interaction with much higher accuracy than ever before. The robustness of our approach can be seen in the validation of known complexes and further improvement such as clever incorporation of increased receptor flexibility and development of accurate peptide conformational ensemble generation will further enhance the range of applications.

## Materials and Methods

### Data set (Table 1 and Supplementary Table S1)

Docking performance and analysis was calibrated and assessed on a benchmark of peptide-protein complexes derived from the PeptiDB database ^50^, filtered according to the following criteria:

(1) *Availability of both the complex and the free receptor structure*, solved by X-ray crystallography (resolution of the complex ≤2.0Å).

(2) *Absence of crystal contacts that could influence the peptide conformation*. In certain cases this further interaction is of biological relevance, leading to receptor multimerization and clustering (e.g. PeptiDB entries involving some of the SH3 domain-peptide interactions, 2AK5 ^51^, and 2J6F ^52^). Since for these cases, obtaining high-resolution models might be challenging without including the symmetry mate, such examples were removed from the dataset.

(3) *Absence of large receptor rearrangement upon peptide binding*. Even though the present implementation of PIPER-FlexPepDock does allow for local conformational changes in the receptor (backbone as well as side chains), accurate modeling of more significant movement of the receptor upon peptide binding (e.g. significant loop movement at the binding interface in PeptiDB entry 1D4T ^53^) require the development of algorithms for efficient modeling of more significant receptor flexibility, which is beyond the scope of the present study.

(4) *Non-redundant dataset*. The criteria above result in a dataset of 42 complexes (**Supplementary Table S1C**) that is very similar to the one used in previous studies by different groups ^12^,^13^,^15^,^16^. To ensure that no bias towards a certain peptide-receptor would be introduced, we extracted a domain non-redundant set (defined by CATH classification ^36^), resulting in the 27 complexes described in this study in detail (**Table 1** and **Supplementary Table S1B**).

The dataset was further divided into two subsets, based on available information about a peptide binding motif (defined in this study based on ELM ^29^, http://elm.eu.org): For the **motif set (**12 complexes) we modeled only the motif part, since it contributes most to binding, and shorter peptides are easier to model. To enable comparison to performance of other protocols, we subsequently also docked the full peptide. For the **non-motif set** (15 complexes), the full peptide was docked.

**Initial calibration set:** For initial calibration, we selected a smaller subset of 9 complexes (**Supplementary Table S1A**). The established protocol was then validated on the remaining complexes, to ensure similar performance and thereby prevent overfitting of the modeling protocol.

### The steps of the PIPER-FlexPepDock protocol

In the following we provide specific details of the different steps of the PIPER-FlexPepDock protocol. For runline commands, see the **Supplementary Materials** section.

### (1) Generation of peptide conformations using Rosetta fragment picker and Rosetta fixbb design

The Rosetta fragment picker ^22^ uses a scoring measure composed of a weighted combination of secondary structure propensity, sequence profile similarity and residue propensities for local regions in the Ramachandran plot ^54^ to map fragments to *vall*, a database of solved high-resolution protein structures. Consequently, the mapped fragments are consistent with the peptide sequence (as defined by a sequence similarity profile generated with PSI-BLAST ^55^) and secondary structure (as predicted using PSIPRED ^56^; even though PSIPRED was shown to perform quite well for shorter sequences ^11^, we use the full protein sequence from which the peptide was derived for PSIPRED and PSIBLAST runs, where available). If the preferred secondary structure is already known (e.g. the alpha helical nuclear receptor box motif) it can be provided instead of PSIPRED predictions. Secondary structural information can also be obtained from experimental techniques such as Circular dichroism (CD) spectroscopy, or approximated by residue Ramachandran local region propensities (derived from statistical analysis of high-resolution protein structure ^57^). The coordinates of the top fifty assigned fragments are extracted from the PDB, and side chains for non-identical residues are modeled using the Rosetta fixbb design algorithm ^58^. The whole process results in an ensemble of 50 fragments for the query peptide sequence.

### (2) Rigid body docking using PIPER

Each of the fifty fragments is globally docked onto the receptor using the PIPER Fourier transform (FFT) docking algorithm, as detailed before ^24^, decomposing the free receptor into independent binding units (either a single domain or repeated, non-decomposable domains; as in Lavi *et al*. ^9^). The calculations are performed for each of 70,000 rotations, and one lowest-energy translation for each rotation is retained. For each fragment docking run the top ranked 250 solutions (total 50x250 = 12500 models) are collected for refinement in the next step (see **Supplementary Figure S3** for a comparison of performance using different numbers of top-ranked solutions).

### Selection of final model from a PIPER simulation

In order to compare performance of a protocol involving only the first PIPER rigid body docking step (in **Table 1**), we selected the final models as reported previously (similar to the PeptiDock implementation ^24^, but without minimization). In short, the models collected are clustered (with radius of 3.5Å C_α_ RMSD), and cluster density is used for ranking and selection of representatives.

### (3) The Rosetta FlexPepDock refinement algorithm

The FlexPepDock Refinement protocol refines all of the peptide’s degrees of freedom (*i.e*. its rigid body orientation as well as backbone dihedral angles), as well as the receptor side chain conformations. Rosetta FlexPepDock refinement was performed as described previously ^10^, with slight changes: (1) *Sampling:* In our present implementation, we also allowed the receptor backbone to move during minimization steps, to allow for slight readjustment upon binding (compare e.g. **Figures 3B** and **3C**). (2) *Scoring:* Rosetta energy function Talaris2014 ^44^ was used. Clustering of models was performed as previously described, using a threshold of 2.0Å ^32^. The top-scoring member of each cluster (according to reweighted score) was selected as the representative member, and clusters were ranked based on the reweighted score of the representative members (as in Raveh *et al*. ^11^).

### Model Evaluation Criteria

For each global docking run the 10 top ranking clusters were selected as prediction and evaluated for quality based on ligand RMSD (L-RMSD), calculated between the native and model peptide backbone atoms after optimal superimposition of the receptor, as done in the CAPRI assessment ^34^,^35^. L-RMSD and other measures, such as Fnat and I-RMSD, were calculated using DockQ ^59^.

### Rosetta release version

The protocol and tests described in this manuscript follow the FlexPepDock protocol, as implemented within the Rosetta weekly release version 2016.20.58704.

### Simulation running time

The processing time for the different stages of the protocol depends on both the length of receptor and the peptide sequence. For example the global docking the carboxy-terminal tail of the ErbB2 Receptor GLDVPV onto the free ERBIN PDZ domain (103 residues) the generation of 50 fragments takes ∼8 CPU minutes over an AMD Sun cluster with 300 cores. For the same complex a single PIPER fragment docking simulation takes ∼2 minutes and a single refinement run of the PIPER docked model takes ∼1 minutes on the same system architecture (∼ 1.5 hours to refine all models).

### Protocol availability

The runline commands are provided in the **Supplementary Material Section**. The Rosetta software is available for free to the academic community. The details regarding downloading and installation is available at **https://www.rosettacommons.org**. PIPER FFT rigid body docking is available as part of the protein-protein docking server ClusPro (PeptiDock at https://peptidock.cluspro.org).

## Acknowledgements

We thank Dr. Barak Raveh for insightful discussions. We also thank Dr. Christina Schindler for providing the pepATTRACT models as reported in Schindler et al. ^13^, and Dr. Mikael Trellet and Prof. Alexandre Bonvin for providing the link for the SBGrid deposited HADDOCK models (<u>https://data.sbgrid.org/dataset/131/</u>) as reported in Trellet et al. ^12^, for the comparison of performance.

## Author contributions

NA conceived the study, designed the project, developed and implemented the protocol, generated and analyzed data, wrote the manuscript; OG, NA, and BX developed the server; KP developed the PIPER peptide docking protocol; DK conceived the study, analyzed the data; OSF conceived the study, designed and coordinated the project, analyzed the data, wrote the manuscript.

## Supplementary Information

### Supplementary Material

**Runline commands**

The runline commands for the different stages are given below:

### 1. Fragment generation using fragment picker

The make_fragments.pl script is used to run PSI-BLAST and PSIPRED to generate the peptide secondary structure and the sequence similarity profile:

~~~
$make_fragments.pl -verbose peptide.fasta
~~~

Rosetta fragment picker is used to assign fragments consistent with the predicted secondary structure and sequence profile to the vall database of high-resolution protein fragments:

~~~
$fragment_picker.linuxgccrelease -database rosetta_database - in:file:vall vall.jul19.2011 -in:file:checkpoint pep_seq.checkpoint - frags:frag_sizes 6 -frags:n_candidates 2000 -frags:n_frags 50 - frags:ss_pred pep_seq.psipred_ss2 psipred -frags:scoring:config psi_L1.cfg -frags:bounded_protocol true
~~~

These assigned fragments are extracted from the Protein Data Bank (including the side-chains). The non-identical residues are mutated using the Rosetta fixbb design protocol:

~~~
$fixbb.linuxgccrelease -database rosetta_database -in:file:s fragment_1.pdb -resfile mutation_resfile -ex1 -ex2 -use_input_sc
~~~

### 2. PIPER Docking

*Step I:* preprocessing the input receptor and fragments using pdbprep.pl and pdbnmd.pl:

~~~
$perl pdbprep.pl receptor.pdb
$perl pdbnmd.pl receptor.pdb ‘?’
~~~

Each of the 50 fragments is similarly processed.

*Step II*: Running PIPER FFT docking:

~~~
$piper.acpharis.omp.20120803 -vv -c1.0 -k4 --msur_k=1.0 --maskr=1.0 -T FFTW_EXHAUSTIVE -R 70000 -t 1 -p atoms.0.0.4.prm.ms.3cap+0.5ace.Hr0rec -f coeffs.0.0.4.motif -r rot70k.0.0.4.prm receptor_nmin.pdb fragment1_nmin.pdb >piper.log
~~~

Each of the fragments is docked onto the receptor.

*Step III*: Top scoring 250 PIPER models are extracted:

~~~
$python apply_ftresult.py -i PIPER_model_ID ft.000.00 rot70k.0.0.4.prm fragment1_nmin.pdb --out-prefix PIPER_model_ID
~~~

Where PIPER_model_ID is an integer value assigned to each transformation for a fragment.

### 3. FlexPepDock Refinement

*Step I:* prepacking the PIPER model

In the PIPER docked model the receptor is replaced with a prepacked receptor. A single prepacked receptor is used.

*Step II: Running refinement*

~~~
$FlexPepDocking.mpi.linuxgccrelease -database rosetta_database - in:file:s PIPER_model_1.pdb -scorefile score.sc -min_receptor_bb - lowres_preoptimize -pep_refine -flexpep_score_only -ex1 -ex2aro - use_input_sc -unboundrot free_receptor.pdb
~~~

Where PIPER_model_1.pdb is the prepacked model.

### 4. Clustering

The top scoring 1% refined models (125) are clustered using the Rosetta cluster application:

~~~
$cluster.linuxgccrelease -in:file:silent decoys.silent top_model_list - in:file:silent_struct_type binary -database $PATH_TO_DB -cluster:radius 2.0 -in:file:fullatom -tags `cat top_refined_list - silent_read_through_errors
~~~

The clusters are ranked based on the top scoring decoys from each clusters, based on reweighted score, and top ranking 10 clusters are selected as putative models.

## Supplementary Figures

**Figure S1.**
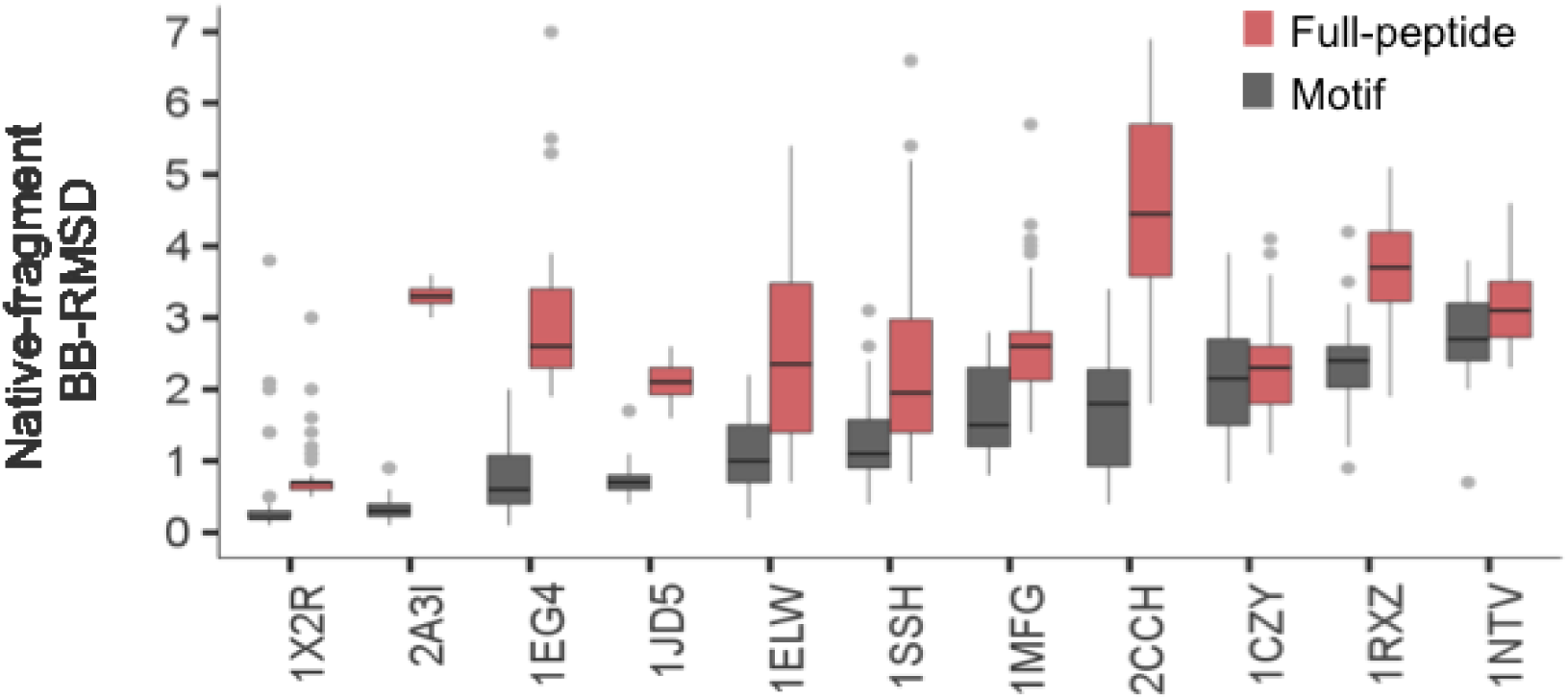
Global peptide docking energy landscapes for the full dataset (accompanies **Figure 3**; provided as separate file). X-axis: L-RMSD, Y-axis: reweighted score. **Top line:** PIPER rigid body docking of peptide fragments onto the unbound receptor structure; **Middle lines:** FlexPepDock refinement of the PIPER docked fragments on the unbound rigid **(second line)** and flexible **(third line)** receptor structure; **Bottom line:** PIPER- FlexPepDock results starting from a bound receptor structure.

**Figure S2.**
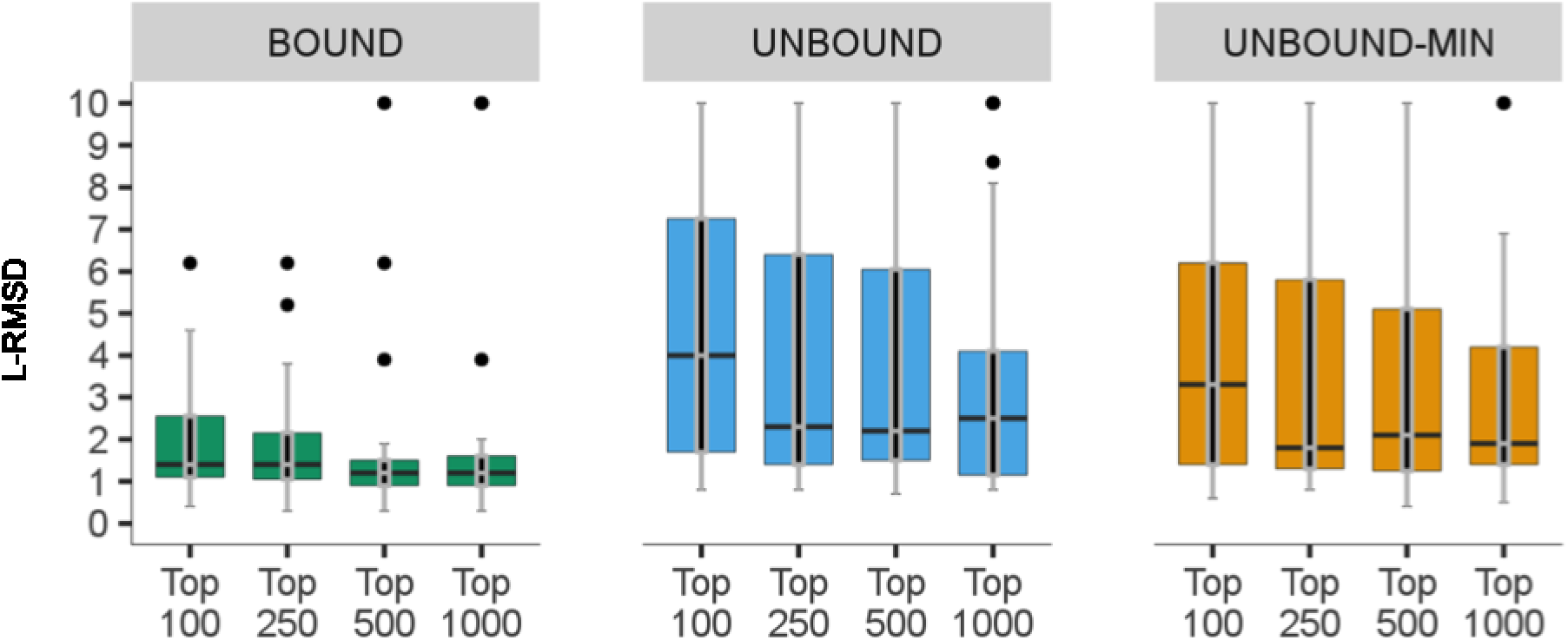
Fragment quality is significantly better for shorter, motif-defined peptide segments (accompanies Figure 2A): Distributions of fragments backbone RMSD values relative to the bound peptide conformations for the motif segments and corresponding full length peptides. The motif set complexes 1JWG and 1TP5 are not added as in these cases the motif covers the whole peptide.

**Figure S3.** The performance of the PIPER-FPD with different number of top PIPER models selected for the refinement stage: Distributions of L-RMSDs of the best models among top 10 ranking clusters for runs using the bound receptor structure (BOUND) and the free receptor structure (UNBOUND & UNBOUND-MIN), the latter including also receptor flexibility in the final refinement step (only the motif region was modeled for the 12 complexes with known motif). The number of PIPER models taken for the FlexPepDock refinement step is shown below each boxplot. Based on these results, we determined a cutoff of 250 models for optimal tradeoff between performance and running time.

## Supplementary Tables

**Table S1.** Details of the datasets of peptide-protein complexes, including modeling results for PIPER-FlexPepDock and other peptide docking protocols (accompanies **Table 1**; provided as separate xls file). **(A)** calibration set (n=9 complexes); **(B)** non-redundant set (n=27 complexes); **(C)** redundant set (n=42 complexes).

**Table S2.** Median fragment-native Backbone-RMSD values for the PeptiDock set complexes obtained using Rosetta fragment picker and the motif-based fragment generation approach (used in PeptiDock ^24^)

| PDB ID |  | Peptide sequence<br>(defined by<br>PeptiDock) | Fragment quality <sup>a</sup> |  |
| --- | --- | --- | --- | --- |
| Complex | Free<br>Receptor |  | Rosetta fragment<br>picker | PeptiDock sequence<br>– motif based<br>fragment picker |
| 1D4T:A | 1D1ZA | TIYAQV | 2.4 | **1.9<sup>b</sup>** |
| 1SSH:A | 1OOTA | PAMPAR | 2.0 | 2.1 |
| 1MFG:A | 2H3LA | LDVPV | 1.4 | 1.4 |
| 2H9M:A | 2H14A | ARTKQ | **2.1** | 2.4 |
| 2FOJ:A | 2FAWA | RAHSS | **1.6** | 2.0 |
| 2HPL:A | 2HPJA | DDLYG | **1.7** | 2.2 |
| 1CZY:A | 1CA4A | PQQATDD | 2.4 | 2.5 |
| 1JD5:A | 1JDA | AIAYF | 1.4 | 1.5 |
| 2VJ0:A | 1B9KA_1 | WVTFE | **1.1** | 2.1 |
| 2VJ0:A | 1B9KA_2 | FEDNF | **2.0** | 2.5 |
| 2C3I:B | 2J2IB | RRRHPS | 2.5 | 2.4 |
| 2CCH:B | 1H1RB | KGRRL | **1.5** | 1.9 |
| 1EG4:A | 1EG3A | RSPPPY | 1.6 | 1.8 |
| 1RXZ:A | 1RWZA | QATLERWF | 2.9 | 2.8 |
| 1ER8:E | 4PAEA | HLLVY | 1.8 | 2.0 |
| 1JWG:A | 1JWFA | DEDLL | 2.1 | **1.6** |

<sup>a</sup> Similarity between fragments and bound peptide conformation: Median backbone RMSD (Å)
<sup>b</sup> In bold: significantly better fragments ( $\Delta \geq 0.3$ Å)

## References

1 Pawson & Nash. Assembly of cell regulatory systems through protein interaction domains. Science 300, 445–452, (2003).

2 Petsalaki & Russell. Peptide-mediated interactions in biological systems: new discoveries and applications. Curr Opin Biotechnol 19, 344–350, (2008).

3 Neduva, Linding, Su-Angrand, Stark, de Masi, Gibson, Lewis, Serrano & Russell. Systematic discovery of new recognition peptides mediating protein interaction networks. PLoS Biol 3, e405, (2005).

4 Vacic, Oldfield, Mohan, Radivojac, Cortese, Uversky & Dunker. Characterization of molecular recognition features, MoRFs, and their binding partners. J Proteome Res 6, 2351–2366, (2007).

5 Gamble, Vajdos, Yoo, Worthylake, Houseweart, Sundquist & Hill. Crystal structure of human cyclophilin A bound to the amino-terminal domain of HIV-1 capsid. Cell 87, 1285–1294, (1996).

6 London, Raveh & Schueler-Furman. Druggable protein-protein interactions-from hot spots to hot segments. Curr Opin Chem Biol 17, 952–959, (2013).

7 Trabuco, Lise, Petsalaki & Russell. PepSite: prediction of peptide-binding sites from protein surfaces. Nucleic Acids Res 40, W423–427, (2012).

8 Saladin, Rey, Thevenet, Zacharias, Moroy & Tuffery. PEP-SiteFinder: a tool for the blind identification of peptide binding sites on protein surfaces. Nucleic Acids Res 42, W221–226, (2014).

9 Lavi, Ngan, Movshovitz-Attias, Bohnuud, Yueh, Beglov, Schueler-Furman & Kozakov. Detection of peptide-binding sites on protein surfaces: the first step toward the modeling and targeting of peptide-mediated interactions. Proteins 81, 2096–2105, (2013).

10 Raveh, London & Schueler-Furman. Sub-angstrom modeling of complexes between flexible peptides and globular proteins. Proteins 78, 2029–2040, (2010).

11 Raveh, London, Zimmerman & Schueler-Furman. Rosetta FlexPepDock ab-initio: simultaneous folding, docking and refinement of peptides onto their receptors. PLoS One 6, e18934, (2011).

12 Trellet, Melquiond & Bonvin. A unified conformational selection and induced fit approach to protein-peptide docking. PLoS One 8, e58769, (2013).

13 Schindler, de Vries & Zacharias. Fully Blind Peptide-Protein Docking with pepATTRACT. Structure 23, 1507–1515, (2015).

14 Ben-Shimon & Niv. AnchorDock: Blind and Flexible Anchor-Driven Peptide Docking. Structure 23, 929–940, (2015).

15 Kurcinski, Jamroz, Blaszczyk, Kolinski & Kmiecik. CABS-dock web server for the flexible docking of peptides to proteins without prior knowledge of the binding site. Nucleic Acids Res 43, W419–424, (2015).

16 Yan, Xu & Zou. Fully Blind Docking at the Atomic Level for Protein-Peptide Complex Structure Prediction. Structure 24, 1842–1853, (2016).

17 Peterson, Roy, Christoffer, Terashi & Kihara. Modeling disordered protein interactions from biophysical principles. PLoS Comput Biol 13, e1005485, (2017).

18 Dominguez, Boelens & Bonvin. HADDOCK: a protein-protein docking approach based on biochemical or biophysical information. J Am Chem Soc 125, 1731–1737, (2003).

19 de Vries, Rey, Schindler, Zacharias & Tuffery. The pepATTRACT web server for blind, large-scale peptide-protein docking. Nucleic Acids Res, (2017).

20 Webb & Sali. Comparative Protein Structure Modeling Using MODELLER. Curr Protoc Bioinformatics 47, 5 6 1–32, (2014).

21 Trott & Olson. AutoDock Vina: improving the speed and accuracy of docking with a new scoring function, efficient optimization, and multithreading. J Comput Chem 31, 455–461, (2010).

22 Gront, Kulp, Vernon, Strauss & Baker. Generalized fragment picking in Rosetta: design, protocols and applications. PLoS One 6, e23294, (2011).

23 Venkatraman, Yang, Sael & Kihara. Protein-protein docking using region-based 3D Zernike descriptors. BMC Bioinformatics 10, 407, (2009).

24 Porter, Bing, Beglov, Bohnuud, Alam, Schueler-Furman & Kozakov. ClusPro PeptiDock: Efficient global docking of peptide recognition motifs using FFT. Bioinformatics ***10.1093/bioinformatics/btx216*.**, (2017).

25 Berman, Westbrook, Feng, Gilliland, Bhat, Weissig, Shindyalov & Bourne. The Protein Data Bank. Nucleic Acids Res 28, 235–242, (2000).

26 Kozakov, Beglov, Bohnuud, Mottarella, Xia, Hall & Vajda. How good is automated protein docking? Proteins 81, 2159–2166, (2013).

27 Brooks, Brooks, Mackerell, Nilsson, Petrella, Roux, Won, Archontis, Bartels, Boresch, Caflisch, Caves, Cui, Dinner, Feig, Fischer, Gao, Hodoscek, Im, Kuczera, Lazaridis, Ma, Ovchinnikov, Paci, Pastor, Post, Pu, Schaefer, Tidor, Venable, Woodcock, Wu, Yang, York & Karplus. CHARMM: the biomolecular simulation program. J Comput Chem 30, 1545–1614, (2009).

28 Kozakov, Brenke, Comeau & Vajda. PIPER: an FFT-based protein docking program with pairwise potentials. Proteins 65, 392–406, (2006).

29 Dinkel, Van Roey, Michael, Kumar, Uyar, Altenberg, Milchevskaya, Schneider, Kuhn, Behrendt, Dahl, Damerell, Diebel, Kalman, Klein, Knudsen, Mader, Merrill, Staudt, Thiel, Welti, Davey, Diella & Gibson. ELM 2016-data update and new functionality of the eukaryotic linear motif resource. Nucleic Acids Res 44, D294–300, (2016).

30 Puntervoll, Linding, Gemund, Chabanis-Davidson, Mattingsdal, Cameron, Martin, Ausiello, Brannetti, Costantini, Ferre, Maselli, Via, Cesareni, Diella, Superti-Furga, Wyrwicz, Ramu, McGuigan, Gudavalli, Letunic, Bork, Rychlewski, Kuster, Helmer-Citterich, Hunter, Aasland & Gibson. ELM server: A new resource for investigating short functional sites in modular eukaryotic proteins. Nucleic Acids Res 31, 3625–3630, (2003).

31 Messih, Lepore & Tramontano. LoopIng: a template-based tool for predicting the structure of protein loops. Bioinformatics 31, 3767–3772, (2015).

32 Gray, Moughon, Wang, Schueler-Furman, Kuhlman, Rohl & Baker. Protein-protein docking with simultaneous optimization of rigid-body displacement and side-chain conformations. J Mol Biol 331, 281–299, (2003).

33 Lensink, Velankar & Wodak. Modeling protein-protein and protein-peptide complexes: CAPRI 6th edition. Proteins 85, 359–377, (2017).

34 Mendez, Leplae, De Maria & Wodak. Assessment of blind predictions of protein-protein interactions: current status of docking methods. Proteins 52, 51–67, (2003).

35 Mendez, Leplae, Lensink & Wodak. Assessment of CAPRI predictions in rounds 3-5 shows progress in docking procedures. Proteins 60, 150–169, (2005).

36 Pearl, Bennett, Bray, Harrison, Martin, Shepherd, Sillitoe, Thornton & Orengo. The CATH database: an extended protein family resource for structural and functional genomics. Nucleic Acids Res 31, 452–455, (2003).

37 Rohl, Strauss, Misura & Baker. Protein structure prediction using Rosetta. Methods Enzymol 383, 66–93, (2004).

38 Park, Lee, Heo & Seok. Protein loop modeling using a new hybrid energy function and its application to modeling in inaccurate structural environments. PLoS One 9, e113811, (2014).

39 Vanhee, Verschueren, Baeten, Stricher, Serrano, Rousseau & Schymkowitz. BriX: a database of protein building blocks for structural analysis, modeling and design. Nucleic Acids Res 39, D435–442, (2011).

40 Li, Suino, Daugherty & Xu. Structural and biochemical mechanisms for the specificity of hormone binding and coactivator assembly by mineralocorticoid receptor. Mol Cell 19, 367–380, (2005).

41 Guhaniyogi, Robinson & Stock. Crystal structures of beryllium fluoride-free and beryllium fluoride-bound CheY in complex with the conserved C-terminal peptide of CheZ reveal dual binding modes specific to CheY conformation. J Mol Biol 359, 624–645, (2006).

42 Todd, Moore, Deivanayagam, Lin, Chattopadhyay, Maki, Wang & Narayana. A structural model for the inhibition of calpain by calpastatin: crystal structures of the native domain VI of calpain and its complexes with calpastatin peptide and a small molecule inhibitor. J Mol Biol 328, 131–146, (2003).

43 Remenyi, Good, Bhattacharyya & Lim. The role of docking interactions in mediating signaling input, output, and discrimination in the yeast MAPK network. Mol Cell 20, 951–962, (2005).

44 Leaver-Fay, O'Meara, Tyka, Jacak, Song, Kellogg, Thompson, Davis, Pache, Lyskov, Gray, Kortemme, Richardson, Havranek, Snoeyink, Baker & Kuhlman. Scientific benchmarks for guiding macromolecular energy function improvement. Methods Enzymol 523, 109–143, (2013).

45 Rego & Koes. 3Dmol.js: molecular visualization with WebGL. Bioinformatics 31, 1322–1324, (2015).

46 Simons, Kooperberg, Huang & Baker. Assembly of protein tertiary structures from fragments with similar local sequences using simulated annealing and Bayesian scoring functions. J Mol Biol 268, 209–225, (1997).

47 Ho & Dill. Folding very short peptides using molecular dynamics. PLoS Comput Biol 2, e27, (2006).

48 Vanhee, Stricher, Baeten, Verschueren, Lenaerts, Serrano, Rousseau & Schymkowitz. Protein-peptide interactions adopt the same structural motifs as monomeric protein folds. Structure 17, 1128–1136, (2009).

49 Kozakov, Li, Hall, Beglov, Zheng, Vakili, Schueler-Furman, Paschalidis, Clore & Vajda. Encounter complexes and dimensionality reduction in protein-protein association. Elife 3, e01370, (2014).

50 London, Movshovitz-Attias & Schueler-Furman. The structural basis of peptide-protein binding strategies. Structure 18, 188–199, (2010).

51 Jozic, Cardenes, Deribe, Moncalian, Hoeller, Groemping, Dikic, Rittinger & Bravo. Cbl promotes clustering of endocytic adaptor proteins. Nat Struct Mol Biol 12, 972–979, (2005).

52 Moncalian, Cardenes, Deribe, Spinola-Amilibia, Dikic & Bravo. Atypical polyproline recognition by the CMS N-terminal Src homology 3 domain. J Biol Chem 281, 38845–38853, (2006).

53 Poy, Yaffe, Sayos, Saxena, Morra, Sumegi, Cantley, Terhorst & Eck. Crystal structures of the XLP protein SAP reveal a class of SH2 domains with extended, phosphotyrosine-independent sequence recognition. Mol Cell 4, 555–561, (1999).

54 Ramachandran, Ramakrishnan & Sasisekharan. Stereochemistry of polypeptide chain configurations. J Mol Biol 7, 95–99, (1963).

55 Altschul, Madden, Schaffer, Zhang, Zhang, Miller & Lipman. Gapped BLAST and PSI-BLAST: a new generation of protein database search programs. Nucleic Acids Res 25, 3389–3402, (1997).

56 Ward, McGuffin, Buxton & Jones. Secondary structure prediction with support vector machines. Bioinformatics 19, 1650–1655, (2003).

57 Berman, Battistuz, Bhat, Bluhm, Bourne, Burkhardt, Feng, Gilliland, Iype, Jain, Fagan, Marvin, Padilla, Ravichandran, Schneider, Thanki, Weissig, Westbrook & Zardecki. The Protein Data Bank. Acta Crystallogr D Biol Crystallogr 58, 899–907, (2002).

58 Kuhlman, Dantas, Ireton, Varani, Stoddard & Baker. Design of a novel globular protein fold with atomic-level accuracy. Science 302, 1364–1368, (2003).

59 Basu & Wallner. DockQ: A Quality Measure for Protein-Protein Docking Models. PLoS One 11, e0161879, (2016).

